# Ferroptosis-like signaling facilitates a potent innate defense against *Plasmodium* infection

**DOI:** 10.1101/257287

**Authors:** Heather S. Kain, Nadia Arang, Elizabeth K. Glennon, Alyse N. Douglass, Denali R. Dudgeon, Jarrod S. Johnson, Alan Aderem, Alexis Kaushansky

## Abstract

The facets of host control during *Plasmodium* liver infection remain largely unknown and conventional innate regulatory pathways are only minimally effective at eliminating parasites^1^^-^^3^. Ferroptosis, a recently described form of iron-dependent cell death that drives accumulation of reactive oxygen species and lipid peroxides, but has not yet been shown to function as an innate immune response^4,5^. Inducing ferroptosis with pharmacologicals or by genetic perturbation of its negative regulators, GPX4 and SLC7a11, dramatically reduces survival of the *Plasmodium* Liver Stage. In contrast, knockdown or knockout of NOX1 or knockdown of TFR1, which are required for ferroptosis, increases the number of Liver Stage parasites. Moreover, we demonstrate that blocking ferroptosis renders parasite-infected hepatocytes resistant to P53-mediated hepatocyte death. Our work establishes that ferroptotic signaling serves to control *Plasmodium* infection in the liver and raises the possibility that ferroptosis operates as an axis of the innate immune system to defend against intracellular pathogens.

---

The expansion and compression of pathogen numbers are a hallmark of infectious lifecycles. For intracellular pathogens, compression can come as the result of host cell death through apoptosis, necrosis, or pyroptosis. In addition to serving as innate defenses, these mechanisms of cell death also inform subsequent adaptive responses (reviewed in ^6^). Other cell death modalities, including ferroptosis^4,5^, have not been explored for their capacity to aid in the elimination of pathogens.

*Plasmodium* parasites, the causative agents of malaria, are first transmitted to mammalian hosts by the bite of an infected *Anopheles* mosquito. After transmission, parasites travel rapidly through the bloodstream to the liver where each parasite infects a hepatocyte to form a Liver Stage (LS)^7,8^. Only after the completion of LS infection do malaria parasites exit the liver, re-enter the bloodstream, infect erythrocytes and initiate symptomatic malaria. Previous evidence suggests that mammalian hosts are, on a cellular level, able to eliminate most malaria parasites prior to blood stage infection. For example, in one study of the rodent malaria *Plasmodium berghei*, the number of sporozoites required to initiate blood stage infection varied from 50 to 10,000 across different strains of naïve mice^9^. These differences, some which have been demonstrated to originate from differences in hepatocyte biology^10^, suggest that variation in host biology can dramatically alter susceptibility to infection.

Conventional innate defenses appear to play only a modest role in curtailing initial infection. Blocking caspase-dependent death in hepatocytes increases LS burden by less than two-fold^3^. The elimination of many classic innate signaling genes such as Toll-like receptor (TLR) 3, TLR4 and the type I interferon receptor have little to no impact on initial LS burden^1,2^. This suggests that other forms of cell death likely contribute to the control of infection.

Necroptosis, or programmed necrosis, has been demonstrated to play a role in innate immunity against a number of pathogens (reviewed in ^6^). Surprisingly, inhibition of necroptosis with the small molecule Necrostatin-1 decreased *Plasmodium yoelii* LS infection of Hepa 1-6 cells (Fig. S1). While this initial experiment does not eliminate the possibility that necroptosis plays a role in host defense against LS infection, cross talk between necroptosis and other forms of cell death have been previously described (reviewed in ^6^), raising the possibility that blocking necroptosis in infected cells sensitizes infected hepatocytes to alternative forms of cell death. We asked if reactive oxygen species (ROS) dependent hepatocyte death might play a role in eliminating LS infection. We infected Hepa 1-6 mouse hepatoma cells with *Plasmodium yoelii* sporozoites and monitored infection and the generation of ROS by flow cytometry. We observed a significant increase in ROS levels in infected hepatocytes (Fig. 1a, Fig. S2). In contrast, we see no significant difference in ROS levels in cells treated with material from uninfected mosquito salivary glands (Fig. S3a).

**Figure 1:**
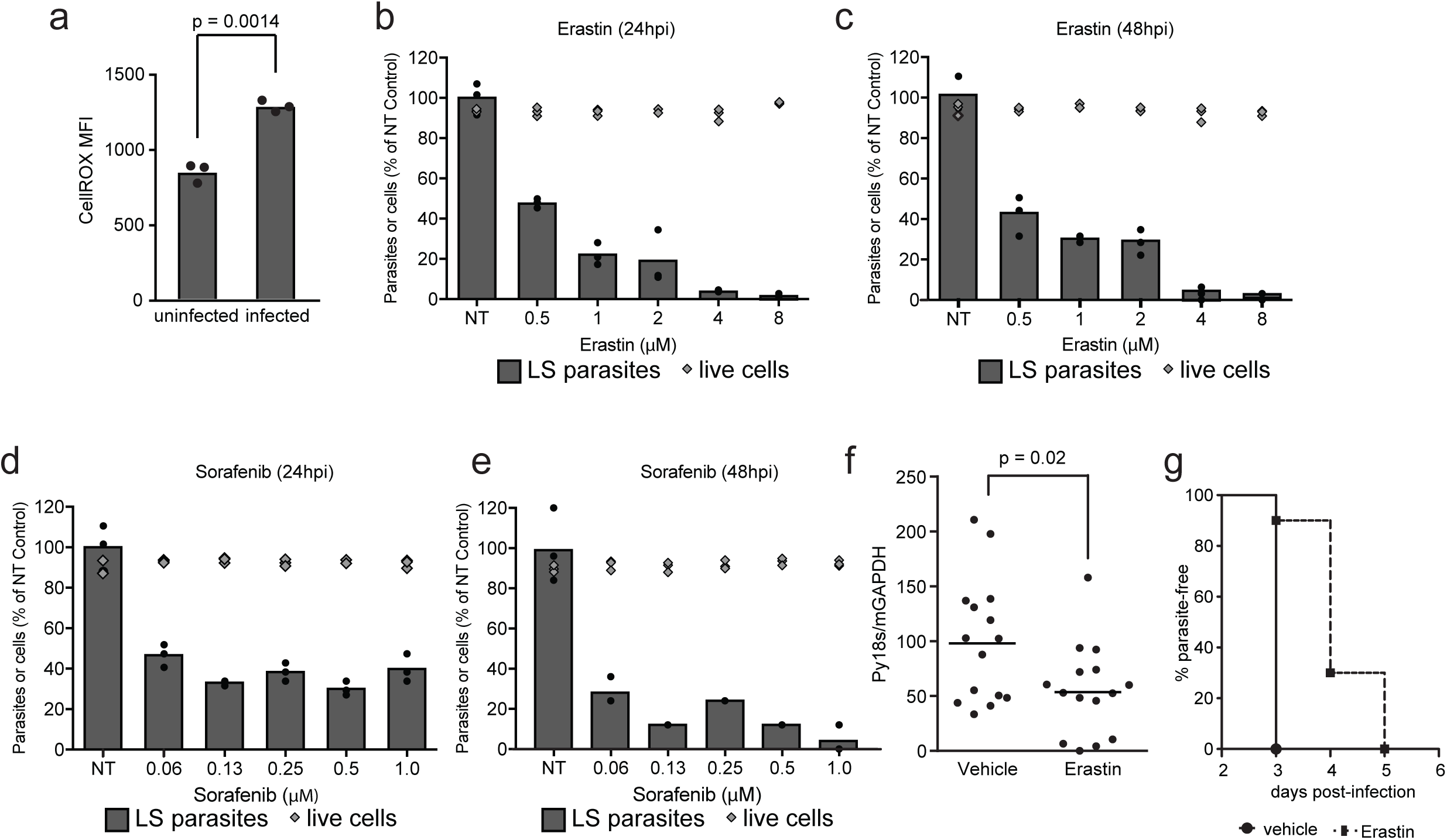
Induction of ferroptosis-like signaling with small molecules eliminates *Plasmodium* LS parasites *in vitro* and *in vivo*. (a) Hepa1-6 cells were infected with 10^5^ *P. yoelii* sporozoites and then evaluated for ROS with a CellROX dye 24 h after infection via flow cytometry. Mean fluorescent intensity (MFI) of signal obtained is presented. P-value was obtained using a Student’s t-test. (b, c) Hepa1-6 cells were infected with 5 x 10^4^ *P. yoelii* sporozoites and treated with Erastin at indicated concentrations 90 min after infection. After (b) 24 hours or (c) 48 hours, LS parasites were visualized by *Py* HSP70 staining and quantified by fluorescent microscopy. Cell death in uninfected cells was quantified by Trypan Blue staining. (d, e) Hepa1-6 cells were infected with *P. yoelii* and treated with Sorafenib at indicated concentrations. After 24 h (d) or 48 h (e), LS parasites were visualized by PyHSP70 staining and quantified by fluorescent microscopy. Cell death in uninfected cells was evaluated by Trypan Blue staining. (f) WT or NOX1^(-/-)^ C57Bl/6 mice were infected via retro-orbital injection with 10^5^ *P. yoelii* sporozoites. Livers were isolated 42 h post-infection and LS burden was quantified by qRT-PCR. P-value was determined by student’s t-test. (g) 10 C57Bl/6 mice were treated with 30 mg/kg Erastin or vehicle control for 4 days. On the second day of treatment, mice were challenged with 1000 *P. yoelii* sporozoites by retro-orbital injection. Beginning on day 3 post-infection, blood was evaluated by giemsa-stained thin smear for the presence of blood stage parasites. In panels a-e, points represent individual technical replicates. Each bar represents the mean of replicates.

Interestingly, previous work has demonstrated that heme oxygenase (HO), an enzyme that promotes the metabolism of free heme and reduces ROS is critical for the promotion of LS infection^11^. Moreover, an iron-deficient diet leads to an increase in LS burden^12^. In combination, these findings led us to question if ferroptosis, a recently described iron and ROS dependent, caspase-independent, non-necroptotic form of cell death, regulates host control of LS infection.

Ferroptosis was originally described as Erastin-mediated cell death^4^ that occurs in rat sarcoma (RAS)-expressing, but not RAS-lacking transformed fibroblasts^13^. More recently, ferroptosis has been demonstrated to occur in other contexts^14^^-^^17^, and it is clear that select cells, but not other cells, are susceptible to death via ferroptosis. To evaluate if LS-infected hepatocytes are sensitized to ferroptosis-like death when compared to uninfected cells, we infected Hepa 1-6 cells with *P. yoelii* sporozoites, and then treated cultures with Erastin beginning 1.5 hr post-infection. We observe a dramatic, dose-dependent reduction in the number of LS parasites at 24 h (Fig. 1b) and 48 h (Fig. 1c) after infection, but no increase in death in uninfected cells across the same range of concentrations. Consistent with the model that ferroptotic-like death is selectively promoted in infected cells, we observe increased levels of ROS in infected cells, but not uninfected cells, in response to Erastin treatment (Fig. S3b). This suggests that *Plasmodium* infection renders infected cells susceptible to ferroptosis, while their uninfected counterparts remain resistant. We observe no statistically significant difference in the size of LS parasites in response to Erastin treatment (Fig. S4). Moreover, we do not observe ferroptosis-related signaling in uninfected Hepa 1-6 cells (Fig. S5a). We hypothesized that *Plasmodium* infected hepatocytes are sensitized to cell death that resembles ferroptosis.

While Erastin treatment is the established method for inducing ferroptosis in culture, drug promiscuity could elicit off-target effects that confound the results. To minimize this possibility, we treated infected cultures with Sorafenib, another inducer of ferroptosis^18^^-^^20^ and observed a similar result (Fig. 1d, e). A third, structurally-unrelated inducer of ferroptosis, RSL3, also dramatically reduces the number of LS parasites without eliminating uninfected Hepa 1-6 cells (Fig. S6). Neither Erastin nor Sorafenib impacted the growth of *Plasmodium*-infected erythrocytes^21^, further minimizing the likelihood that these drugs directly target parasite pathways (Fig. S5b).

To evaluate if a similar sensitivity to ferroptotic signaling exists *in vivo*, we treated C57Bl/6 mice with 30 mg/kg Erastin for three days. On the second day of treatment, we infected mice with 10^5^ *P. yoelii* sporozoites by intravenous injection. We observed a significant decrease in LS burden 44 h post-infection (Fig. 1f). Next, we treated mice with 30mg/kg Erastin for 4 days and challenged them with 1000 *P. yoelii* sporozoites on the second day of treatment. We reasoned that any decrease in LS infection would result in a delay in the onset of blood stage infection and monitored the onset of blood stage patency by thin smear in the mice. We observed a 1-2 day delay in the onset of blood stage infection (Fig. 1g). These data support the hypothesis that LS-infected hepatocytes are sensitized to ferroptosis compared to uninfected hepatocytes both *in vivo* and *in vitro*.

Glutathione peroxidase 4 (GPX4) and the cysteine/glutamate transporter Solute carrier family 7 member 11 (SLC7a11/xCT) have been described as negative regulators of ferroptosis^14,22^. After knockdown of the negative regulators of ferroptosis, GPX4 and SLC7a11, in Hepa 1-6 cells (Fig. 2a, b), we observed a significant decrease in the number of LS parasites (Fig. 2c). Moreover, the effect of Erastin is completely ablated in cells that are transduced with sgRNAs against SLC7a11, suggesting that Erastin acts in a SLC7a11-dependent manner (Fig. S7b). In contrast, NADPH oxidase 1 (NOX1)^4^ and iron transport through the transferrin receptor protein 1 (TFR1)^23^ are positive regulators of the process. Accordingly, the knockdown of TFR1 or NOX1 substantially increased the number of LS parasites (Fig. 2d). To examine whether this pathway is relevant *in vivo*, and to evaluate the impact of complete NOX1 knockout, we compared infection in WT and NOX1^(-/-)^ C57Bl/6 mice. NOX1^(-/-)^ mice exhibited a dramatic increase in LS burden (Fig. 2e). For comparison, eliminating apoptosis, multiple toll-like receptors, or type I interferon signaling at most doubles LS parasite burden^1^^-^^3^ in naïve mice. Strikingly, NOX1^(-/-)^ mice exhibited an average of 73 times more LS burden than WT animals. This suggests that NOX1-mediated signaling constitutes a major axis of host control of LS infection.

**Figure 2:**
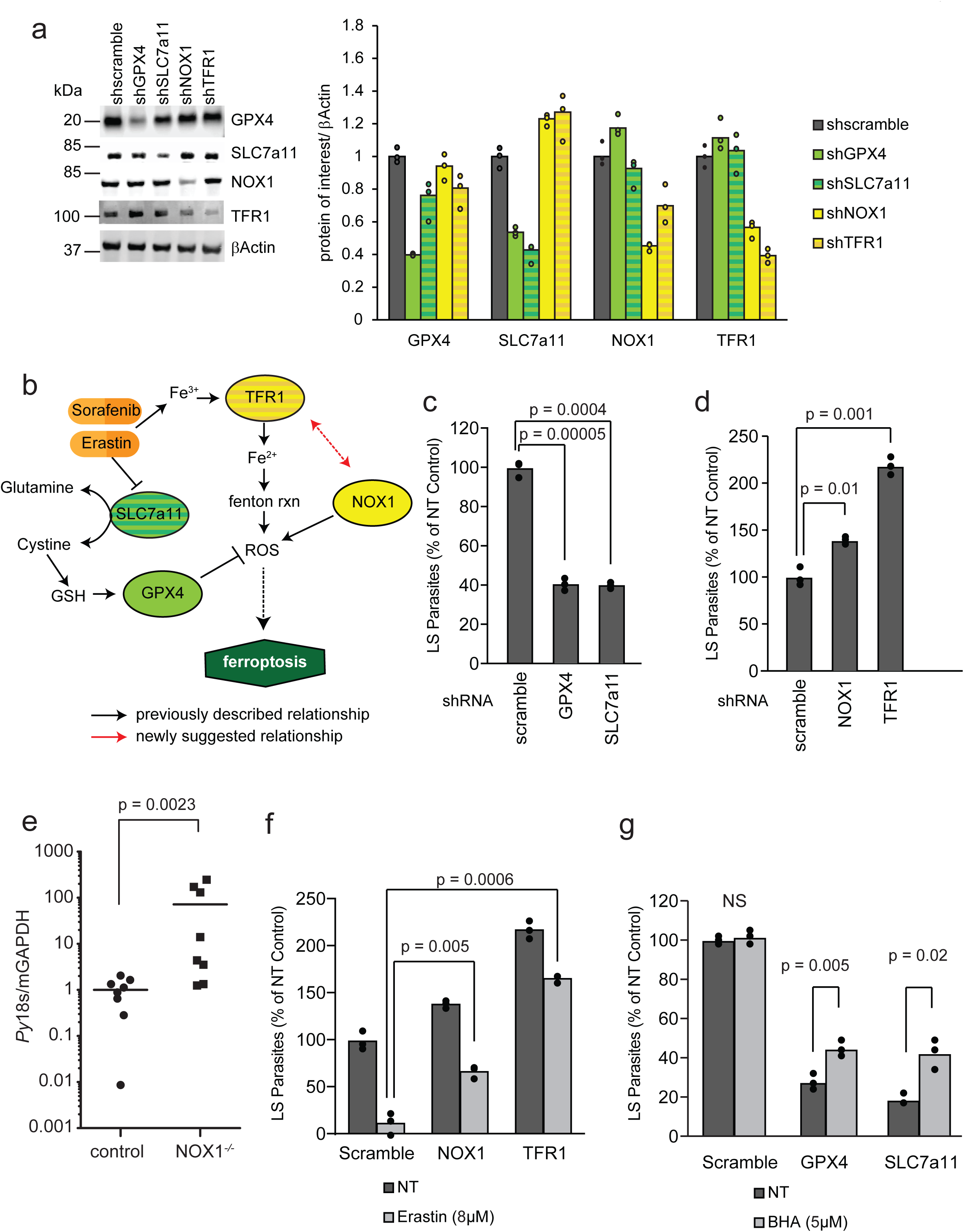
Ferroptosis-related signaling is a potent regulator of LS *Plasmodium* infection. (a) Hepa1-6 cells were transduced with 2-4 lentivirus expressing shRNAs against each gene of interest. Cellular lysates were isolated after lentivirus transduction and selection with puromycin and western immunoblots were performed against the indicated targets. Signal was normalized to βActin. (b) Schematic of ferroptotic signaling as reported in the literature. (c, d) Hepa1-6 cells were transduced with lentivirus expressing shRNAs against (c) GPX4, SLC7a11 or (d) NOX1 and TFR1 as well as a non-targeting shRNA control (scramble). 1.5×10^5^ cells were infected with 5×10^4^ *P. yoelii* sporozoites and quantified by microscopy. (e) WT or NOX1^(-/-)^ C57Bl/6 mice were infected via retro-orbital injection with 10^5^ *P. yoelii* sporozoites. Livers were isolated 42 h post-infection and LS burden was quantified by qRT-PCR. (f) Hepa 1-6 cells were transduced with lentivirus expressing shRNAs against a scramble control, GPX4 or SLC7a11. 1.5×10^5^ cells were infected with 5×10^4^ *P. yoelii* sporozoites. 90 minutes post-infection, cells were treated with butylated hydroxyanisole (BHA) or left untreated. Parasites were visualized 24 h post-infection by Hsp70 staining and quantified by microscopy. (g) Hepa 1-6 cells were transduced by lentivirus expressing shRNAs against a scramble control, NOX1 or TFR1. 1.5×10^5^ cells were infected with 5×10^4^ *P. yoelii* sporozoites. 90 minutes post-infection, cells were treated with Erastin or a DMSO control. Parasites were visualized by *Py* Hsp70 staining 24 h post-infection and quantified by microscopy. In all panels, points represent individual analytical replicates. Each bar represents the mean of replicates. P-values were obtained using a Student’s t-test.

We next asked if the activity of Erastin on LS-infected hepatocytes was specific to ferroptotic signaling by observing the effects of Erastin treatment in the context of reduced NOX1 or TFR1. The knockdown of either NOX1 or TFR1 significantly reduced the efficacy of Erastin (Fig. 2f). Taking a complementary approach, we employed the small molecule butylated hydroxyanisole (BHA), which scavenges ROS and mitigates ferroptosis. The addition of BHA to infections performed after SLC7a11 or GPX4 knockdown partially rescued LS infection (Fig. 2g). Thus, both pharmacological and genetic perturbations to signal transduction cascades that have previously been attributed to ferroptosis-like death strongly impact LS parasite levels.

Ferroptosis has been demonstrated to play a role in multiple aberrant signaling states such as cancer^14,15^, neurodegeneration,^16^ and hemorrhagic stroke^17^. The induction of ferroptosis has also been linked to the capacity of P53 to act as a potent tumor suppressor independent of its initiation of apoptosis, cell cycle arrest or senescence^24^. We have previously demonstrated that elevated levels of P53 reduces LS burden^25^. An increase in P53 protein levels can either be achieved by genetic means^25^ or by the addition of the small molecule Nutlin-3^26^. Nutlin-3 binds the E3 ubiquitin ligase MDM-2, which under normal conditions acts to degrade P53^26^. Thus, the addition of Nutlin-3 dissociates MDM-2 and P53, which increases levels of P53, and in turn, reduces LS burden^25,27^. Interestingly, this reduction in parasite burden cannot be reversed by blocking apoptosis ^27^, nor is it likely associated with P53’s capacity to arrest the cell cycle^28,29^. To evaluate if elevated P53 levels induce ferroptotic signaling, we treated Hepa1-6 cells with 10 μM Nutlin-3 for 24 h and observed suppression of SLC7a11 and GPX4 (Fig. 3a, b). We next asked if the anti-parasitic activity of Nutlin-3 required ferroptotic signaling. When we compare the efficacy of Nutlin-3 in cells transduced with a control shRNA to those with decreased NOX1 or TFR1, we observe a complete loss of susceptibility of LS-infected cells to Nutlin-3 treatment (Fig. 3c). Similarly, when the ferroptotic pathway was blocked using BHA or ferrostatin-1, infected Hepa1-6 cells lost their susceptibility to Nutlin-3 (Fig. 3d).

**Figure 3:**
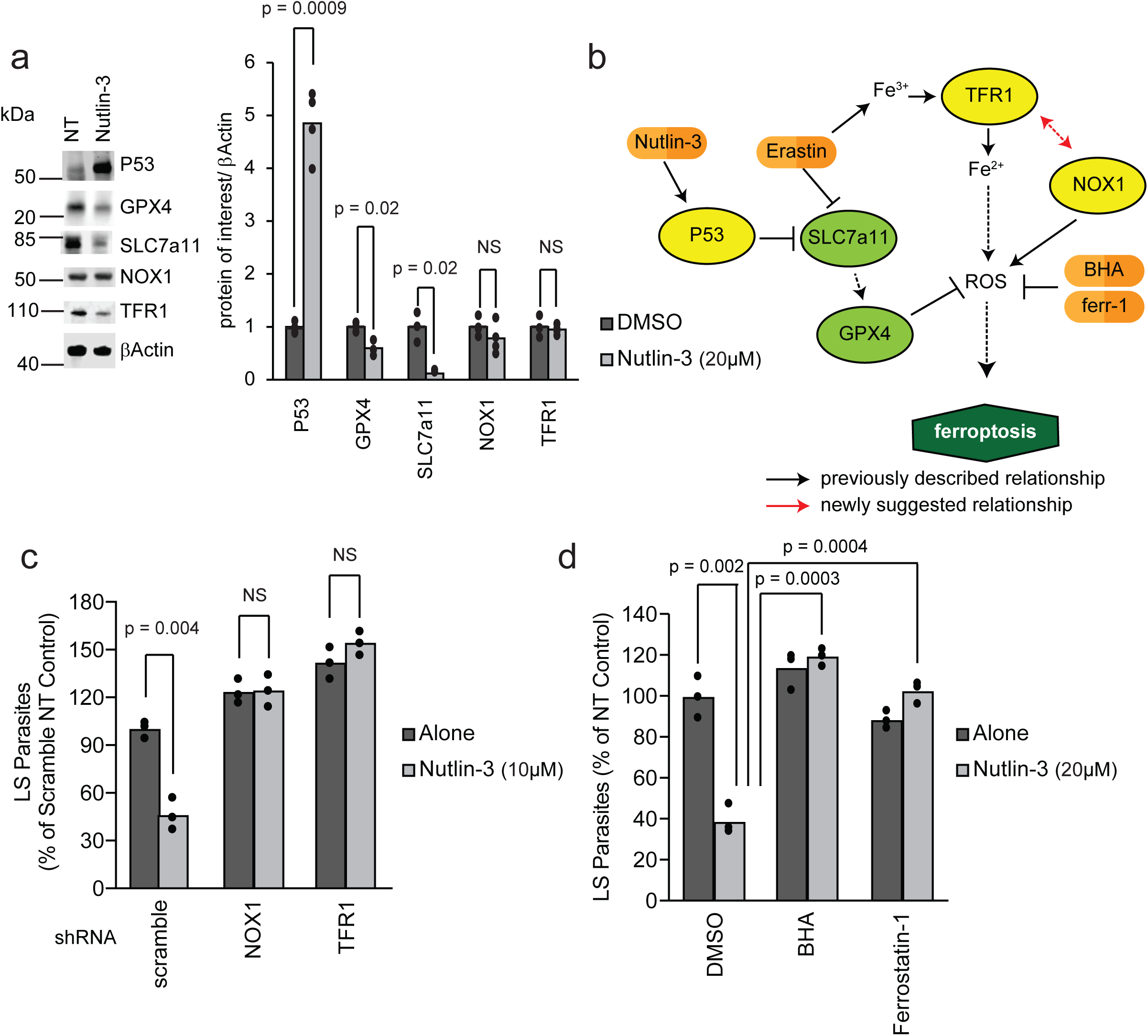
Ferroptotic signaling is responsible for P53-mediated elimination of LS-infected hepatocytes. (a) 1.5×10^5^ Hepa1-6 cells were treated with 10 μM Nutlin-3. Cellular lysates were isolated 24 h after treatment and immunoblots against the indicated targets were performed. Signal was quantified using Image Studio software and normalized to βActin. (b) Schematic of ferroptotic signaling induced by P53. (c) 1.5×10^5^ Hepa 1-6 cells were transduced with lentivirus expressing shRNAs against a scramble control, NOX1 or TFR1 and infected with 5×10^4^ *P. yoelii* sporozoites. 90 minutes post-infection, cells were treated with 10 μM Nutlin-3 or a DMSO control. Parasites were visualized 24 h post-infection by Hsp70 staining and quantified by microscopy. (d) 1.5×10^5^ Hepa1-6 cells were infected with 5×10^4^ *P. yoelii* sporozoites. 90 minutes post-infection, cells were treated with a DMSO control, 20 μM Nutlin-3, 5 μM BHA or 300 nM ferrostatin-1 as indicated. Parasites were visualized by *Py* HSP70 staining 24 h post-infection and quantified by microscopy. In all panels, points represent individual analytical replicates. Each bar represents the mean of replicates. P-values were obtained using a Student’s t-test.

The role of P53 in curtailing malaria infection is not dependent on its conventional roles in apoptosis^27^ or cell cycle arrest^28^ but instead, we propose, depends on its ability to initiate ferroptosis. As such, our work further establishes non-canonical activities of P53 as critical for the control of aberrant signaling states. Moreover, we demonstrate that ferroptotic signaling can act as a potent innate defense against intracellular infection and that infection can be further curtailed by inducing ferroptosis with small molecules or by genetic perturbations of the signaling cascade. Interestingly, there is some evidence that the Erastin-induced pathway that controls LS infection differs from canonical ferroptosis as ROS scavenging does not eliminate the effect of Erastin treatment yet does negate the impact of Nutlin-3 (Fig. S7). As such, the full complement of molecular players that contribute to the innate defense through ferroptosis-like death remain to be characterized. Future work will likely establish whether the reach of this signal transduction cascade extends beyond LS malaria infection and operates to control other pathogens.

The nature of pathogen clearance likely has consequences beyond the capacity to control infection during the exposure of naïve individuals. It was recently described in an elegant set of experiments that transcriptomic responses along with the cell phenotypic responses during cell death could inform adaptive responses to infection^30^. This finding is particularly relevant as evidence mounts that the heterogeneity across cells might be much greater than originally appreciated^31^^-^^33^. Indeed, the innate and adaptive systems that control infection may engage a broader range of molecular players, including ferroptosis, than we have traditionally incorporated into our understanding of immunity.

## ACKNOWLEDGEMENTS

We thank Photini Sinnis and Fidel Zavala for the *P. yoelii* Hsp70 antisera, Stefan Kappe for the gift of the *P. falciparum* GFP-Luciferase strain and the Center for Infectious Disease Research vivarium staff for their work with mice. All work was done according to IACUC procedures and protocols. The authors declare no financial conflicts of interest. This work was funded by R01GM101183 and K99/R00AI111785 to AK and R01AI032972 and R01AI025032 to AA. The data that support the findings of this study are available from the corresponding author upon reasonable request.

## AUTHORS CONTRIBUTIONS

HSK, JSJ, AA and AK designed the experiments, HSK, NA, EKG, AND, DRD, JSJ and performed the experiments, HSK and AK wrote the paper with contributions from all other authors.

## References

1. Miller, J.L., Sack, B.K., Baldwin, M., Vaughan, A.M. & Kappe, S.H. Interferon-mediated innate immune responses against malaria parasite liver stages. Cell reports 7, 436–447 (2014).

2. Liehl, P., et al. Host-cell sensors for Plasmodium activate innate immunity against liver-stage infection. Nature medicine 20, 47–53 (2014).

3. Kaushansky, A., et al. Malaria parasite liver stages render host hepatocytes susceptible to mitochondria-initiated apoptosis. Cell death & disease 4, e762 (2013).

4. Dixon, S.J., et al. Ferroptosis: an iron-dependent form of nonapoptotic cell death. Cell 149, 1060–1072 (2012).

5. Xie, Y., et al. Ferroptosis: process and function. Cell death and differentiation 23, 369–379 (2016).

6. Jorgensen, I., Rayamajhi, M. & Miao, E.A. Programmed cell death as a defence against infection. Nature reviews. Immunology 17, 151–164 (2017).

7. Kaushansky, A. & Kappe, S.H. Selection and refinement: the malaria parasite’s infection and exploitation of host hepatocytes. Current opinion in microbiology 26, 71–78 (2015).

8. Lindner, S.E., Miller, J.L. & Kappe, S.H. Malaria parasite pre-erythrocytic infection: preparation meets opportunity. Cell Microbiol 14, 316–324 (2012).

9. Scheller, L.F., Wirtz, R.A. & Azad, A.F. Susceptibility of different strains of mice to hepatic infection with Plasmodium berghei. Infection and immunity 62, 4844–4847 (1994).

10. Kaushansky, A., et al. Susceptibility to Plasmodium yoelii preerythrocytic infection in BALB/c substrains is determined at the point of hepatocyte invasion. Infection and immunity 83, 39–47 (2015).

11. Epiphanio, S., et al. Heme oxygenase-1 is an anti-inflammatory host factor that promotes murine plasmodium liver infection. Cell Host Microbe 3, 331–338 (2008).

12. Goma, J., Renia, L., Miltgen, F. & Mazier, D. Effects of iron deficiency on the hepatic development of Plasmodium yoelii. Parasite 2, 351–356 (1995).

13. Yagoda, N., et al. RAS-RAF-MEK-dependent oxidative cell death involving voltage-dependent anion channels. Nature 447, 864–868 (2007).

14. Yang, W.S., et al. Regulation of ferroptotic cancer cell death by GPX4. Cell 156, 317–331 (2014).

15. Sun, X., et al. HSPB1 as a novel regulator of ferroptotic cancer cell death. Oncogene 34, 5617–5625 (2015).

16. Hambright, W.S., Fonseca, R.S., Chen, L., Na, R. & Ran, Q. Ablation of ferroptosis regulator glutathione peroxidase 4 in forebrain neurons promotes cognitive impairment and neurodegeneration. Redox biology 12, 8–17 (2017).

17. Zille, M., et al. Neuronal Death After Hemorrhagic Stroke In Vitro and In Vivo Shares Features of Ferroptosis and Necroptosis. Stroke 48, 1033–1043 (2017).

18. Lachaier, E., et al. Sorafenib induces ferroptosis in human cancer cell lines originating from different solid tumors. Anticancer research 34, 6417–6422 (2014).

19. Louandre, C., et al. Iron-dependent cell death of hepatocellular carcinoma cells exposed to sorafenib. International journal of cancer 133, 1732–1742 (2013).

20. Houessinon, A., et al. Metallothionein-1 as a biomarker of altered redox metabolism in hepatocellular carcinoma cells exposed to sorafenib. Molecular cancer 15, 38 (2016).

21. Arang, N., et al. Identifying host regulators and inhibitors of liver stage malaria infection using kinase activity profiles. Nat Commun 8, 1232 (2017).

22. Dixon, S.J., et al. Pharmacological inhibition of cystine-glutamate exchange induces endoplasmic reticulum stress and ferroptosis. eLife 3, e02523 (2014).

23. Gao, M., Monian, P., Quadri, N., Ramasamy, R. & Jiang, X. Glutaminolysis and Transferrin Regulate Ferroptosis. Molecular cell 59, 298–308 (2015).

24. Jiang, L., et al. Ferroptosis as a p53-mediated activity during tumour suppression. Nature 520, 57–62 (2015).

25. Kaushansky, A., et al. Suppression of host p53 is critical for Plasmodium liver-stage infection. Cell reports 3, 630–637 (2013).

26. Vassilev, L.T., et al. In vivo activation of the p53 pathway by small-molecule antagonists of MDM2. Science 303, 844–848 (2004).

27. Douglass, A.N., et al. Host-based Prophylaxis Successfully Targets Liver Stage Malaria Parasites. Molecular therapy: the journal of the American Society of Gene Therapy 23, 857–865 (2015).

28. Austin, L.S., Kaushansky, A. & Kappe, S.H. Susceptibility to Plasmodium liver stage infection is altered by hepatocyte polyploidy. Cell Microbiol 16, 784–795 (2014).

29. Hanson, K.K., March, S., Ng, S., Bhatia, S.N. & Mota, M.M. In vitro alterations do not reflect a requirement for host cell cycle progression during Plasmodium liver stage infection. Eukaryotic cell 14, 96–103 (2015).

30. Yatim, N., et al. RIPK1 and NF-kappaB signaling in dying cells determines cross-priming of CD8(+) T cells. Science 350, 328–334 (2015).

31. Shalek, A.K., et al. Single-cell RNA-seq reveals dynamic paracrine control of cellular variation. Nature 510, 363–369 (2014).

32. Avraham, R., et al. Pathogen Cell-to-Cell Variability Drives Heterogeneity in Host Immune Responses. Cell 162, 1309–1321 (2015).

33. Gaublomme, J.T., et al. Single-Cell Genomics Unveils Critical Regulators of Th17 Cell Pathogenicity. Cell 163, 1400–1412 (2015).

